# Conjugative delivery of CRISPR-Cas9 for the selective depletion of antibiotic-resistant enterococci

**DOI:** 10.1101/678573

**Authors:** Marinelle Rodrigues, Sara W. McBride, Karthik Hullahalli, Kelli L. Palmer, Breck A. Duerkop

## Abstract

The innovation of new therapies to combat multidrug-resistant (MDR) bacteria is being outpaced by the continued rise of MDR bacterial infections. Of particular concern are hospital-acquired infections (HAIs) recalcitrant to antibiotic therapies. The Gram-positive intestinal pathobiont *Enterococcus faecalis* is associated with HAIs and some strains are MDR. Therefore, novel strategies to control *E. faecalis* populations are needed. We previously characterized an *E. faecalis* Type II CRISPR-Cas system and demonstrated its utility in the sequence-specific removal of antibiotic resistance determinants. Here we present work describing the adaption of this CRISPR-Cas system into a constitutively expressed module encoded on a pheromone-responsive conjugative plasmid that efficiently transfers to *E. faecalis* for the selective removal of antibiotic resistance genes. Using *in vitro* competition assays, we show that these CRISPR-Cas-encoding delivery plasmids, or CRISPR-Cas antimicrobials, can reduce the occurrence of antibiotic resistance in enterococcal populations in a sequence-specific manner. Furthermore, we demonstrate that deployment of CRISPR-Cas antimicrobials in the murine intestine reduces the occurrence of antibiotic-resistant *E. faecalis* by several orders of magnitude. Finally, we show that *E. faecalis* donor strains harboring CRISPR-Cas antimicrobials are immune to uptake of antibiotic resistance determinants *in vivo*. Our results demonstrate that conjugative delivery of CRISPR-Cas antimicrobials may be adaptable for future deployment from probiotic bacteria for exact targeting of defined MDR bacteria or for precision engineering of polymicrobial communities in the mammalian intestine.

**Importance:** CRISPR-Cas nucleic acid targeting systems hold promise for the amelioration of multidrug-resistant enterococci, yet the utility of such tools in the context of the intestinal environment where enterococci reside is understudied. We describe the development of a CRISPR-Cas antimicrobial, deployed on a conjugative plasmid, for the targeted removal of antibiotic resistance genes from intestinal *Enterococcus faecalis*. We demonstrate that CRISPR-Cas targeting reduces antibiotic resistance of *E. faecalis* by several orders of magnitude in the intestine. Although barriers exist that influence the penetrance of the conjugative CRISPR-Cas antimicrobial among target recipient *E. faecalis* cells, the removal of antibiotic resistance genes in *E. faecalis* upon uptake of the CRISPR-Cas antimicrobial is absolute. In addition, cells that obtain the CRISPR-Cas antimicrobial are immunized against the acquisition of new antibiotic resistance genes. This study suggests a potential path toward plasmid based CRISPR-Cas therapies in the intestine.

## Introduction

Disruption of the intestinal microbiota can predispose individuals to infection by opportunistic pathogens (1, 2). Antibiotics facilitate such disturbances leading to the development of hospital-acquired infections (HAIs) (3, 4). *Enterococcus faecalis*, a normal constituent of the healthy human intestinal microbiota and historically used in probiotics and during food fermentation, is now a leading cause of HAIs (5–7). *E. faecalis* is an adept opportunist that can proliferate in the dysbiotic intestine following antibiotic perturbation (8). Antibiotic-resistant clinical isolates of *E. faecalis* typically possess expanded genomes compared to susceptible isolates due to the acquisition of mobile genetic elements (9, 10), and patients colonized with these multidrug-resistant (MDR) *E. faecalis* are at increased risk of acquiring bloodstream infections (11). Thus, there is a need for novel strategies to decolonize high-risk individuals of MDR *E. faecalis* (12).

CRISPR-Cas systems are protective barriers in prokaryotes that function in the adaptive immunity to mobile genetic elements (13–15). The well-studied Type II CRISPR-Cas system consists of a DNA endonuclease (Cas9) that uses a programmable RNA guide to cleave a double-stranded DNA target sequence (16). Double-stranded DNA breaks can be lethal to bacteria (17), therefore deploying Type II CRISPR-Cas as a sequence-specific antimicrobial is an attractive alternative to conventional antibiotic therapy. Previous studies have explored this concept using phages to deliver engineered CRISPR systems targeting specific genes (18–21). Although effective, this method of delivery is likely to be challenging for many species of bacteria including *E. faecalis* due to phage receptor and other cell wall mutations that arise readily in response to lytic phage infection (22–25).

We previously demonstrated the utility of CRISPR-Cas9 as a sequence-specific *in vitro* antimicrobial in *E. faecalis* by delivering CRISPR guide sequences on mobilizable cloning vectors into *E. faecalis* cells that chromosomally encode *cas9* (26). However, since MDR *E. faecalis* strains lack complete CRISPR-Cas systems, and specifically *cas9* (27), efficient delivery of the entire CRISPR-Cas9 machinery to MDR *E. faecalis* would likely be required. An alternative method for the delivery of CRISPR-Cas is the use of pheromone-responsive plasmids (PRPs), which naturally achieve high rates of conjugation, have a narrow host range limited to *E. faecalis*, are capable of comprehensively infiltrating *E. faecalis* populations, and can disseminate within intestinal *E. faecalis* in the absence of antibiotic selection (28–31). The transmission of PRPs is tightly regulated, so the deployment of a PRP CRISPR-Cas antimicrobial could be conditionally tuned for precision targeting and delivery (32).

PRPs respond to pheromones (short peptides) that are secreted by plasmid-free *E. faecalis* cells (33). When donor cells detect excess pheromones, indicating the presence of plasmid-free cells in the vicinity (34), they synthesize aggregation substance which facilitates contact between plasmid-bearing and plasmid-free cells (35). This enables the rapid and efficient transfer of PRPs from donor to recipient. In addition, many PRPs encode accessory genes such as antibiotic resistance and bacteriocins, small ribosomally synthesized antimicrobial proteins that target diverse bacteria and for which producing strains must encode immunity genes (36–38). Previous work demonstrated that the PRP pPD1 conjugated at high rates in the intestines of mice, and pPD1 transfer to plasmid-free *E. faecalis* was beneficial since recipients that failed to acquire the plasmid were killed by the pPD1 encoded bac-21 bacteriocin (29).

We recently discovered that *E. faecalis* cells possessing self-targeting CRISPR-Cas systems are able to transiently tolerate this self-targeting, albeit at a fitness cost, and that antibiotic selection determines the outcome of this conflict by selecting for CRISPR mutants or loss of the CRISPR-targeted gene (26, 39). We observed CRISPR-Cas self-targeting lethality when the expression of *cas9* was increased using a constitutive promoter (40). In this study, we engineered pPD1 with a complete, constitutively expressed CRISPR-Cas9 targeting cassette, specific for the enterococcal antibiotic resistance genes *ermB* (encoding erythromycin resistance) and *tetM* (encoding tetracycline resistance). We chose pPD1 as it lacks natively encoded antibiotic resistance yet it harbors the fitness-enhancing bac-21 bacteriocin (41, 42). Using this engineered PRP, we successfully depleted tetracycline and erythromycin resistance from *E. faecalis* populations *in vitro*. Crucially, this approach worked in the absence of externally applied selection for pPD1 in donor strains, showing practicality for its usage as a potential antimicrobial. Using an *in vivo* intestinal colonization model, we show that these constructs are conjugated to erythromycin-resistant recipient cells *in vivo*. Donors carrying the engineered PRP targeting *ermB* significantly reduced the prevalence of erythromycin-resistant intestinal *E. faecalis*, supporting the utility of the engineered PRP in mitigating MDR *E. faecalis*. Furthermore, we show that these donors are immune to the uptake of *ermB*-encoding PRPs harbored by recipient *E. faecalis* cells. This work is a first step in utilizing conjugative elements for specific delivery of CRISPR-Cas antimicrobials to precisely target antibiotic-resistant bacteria in the intestinal microbiota.

## Results

### Delivery of CRISPR-Cas9 effectively removes antibiotic resistance *in vitro* by targeting plasmid-borne resistance genes

To assess the ability of CRISPR-Cas9 to target and remove antibiotic resistance in *E. faecalis*, we modified the PRP pPD1 by introducing the *cas9* gene and a CRISPR guide RNA under the control of the constitutive *bacA* promoter (Fig. 1A and S1). To facilitate detection of the plasmid we included a chloramphenicol resistance marker (*cat*) for selection. We generated two pPD1 derivatives, referred to as pKH88[sp-*tetM*] and pKH88[sp-*ermB*], which carry guide RNAs that target the enterococcal *tetM* and *ermB* genes conferring tetracycline and erythromycin resistance, respectively. These plasmids retain pheromone response functions and bac-21 bacteriocin production (Fig. 1A).

**Figure 1.**
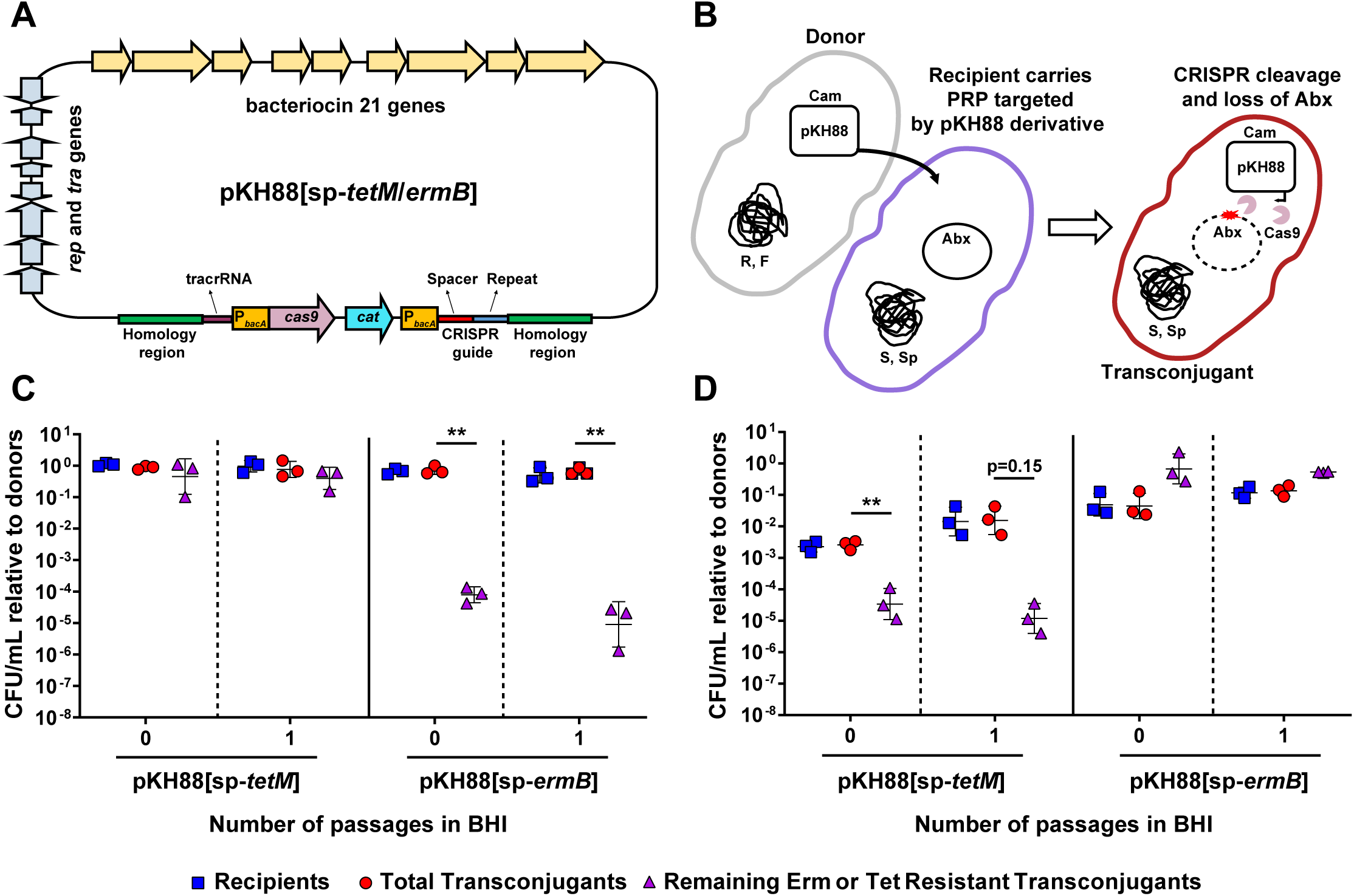
CRISPR-Cas engineered conjugative pPD1 derivative plasmids target and eliminate antibiotic resistance genes in *in vitro* co-culture. **(A)** Schematic of the pPD1 derivative pKH88. pKH88 encodes *cas9* and a CRISPR guide RNA under the control of the constitutive enterococcal *bacA* promoter. The guide RNA consists of a CRISPR repeat region and a unique spacer sequence with complementarity to the *ermB* gene (pKH88[sp-*ermB*]) of pAM771 or the *tetM* gene (pKH88[sp-*tetM*]) of pCF10. pKH88 derivatives encode genes necessary for conjugative transfer, chloramphenicol resistance (*cat*) and biosynthesis of the bacteriocin bac-21. **(B)** Cartoon depicting the delivery of pKH88 derivatives to target recipient *E. faecalis* cells. Abundance of erythromycin **(C)** or tetracycline **(D)** resistant *E. faecalis* OG1SSp recipients following acquisition of pKH88[sp-*ermB*] or pKH88[sp-*tetM*]. Removal of erythromycin and tetracycline resistance from the transconjugant population depends on the introduction of a cognate spacer targeting pKH88 derivative. Markers; R – rifampicin, F – fusidic acid, Cam – chloramphenicol. Abx – antibiotic resistance. **p=0.006 (for passage 0 in C) **p=0.003 (for passage 1 in C), **p=0.009 (for passage 0 in D).

PRPs are vehicles for the transmission of antibiotic resistance genes in *E. faecalis* populations (30), thus we first assessed the ability of our engineered pPD1 derivatives to eliminate antibiotic resistance genes present on other PRPs. We reasoned that bacteriocin production by pKH88 derivatives would promote maintenance of these plasmids in enterococcal populations. The donor strain used in these experiments was *E. faecalis* CK135 and the recipient was *E. faecalis* OG1SSp possessing either pAM771 which carries the erythromycin resistance gene *ermB* or pCF10 which carries the tetracycline resistance gene *tetM* (Fig. 1B). Both CK135 and OG1SSp are isogenic derivatives of the human oral commensal isolate OG1 (43). We mixed donors and recipients and plated them on Brain Heart Infusion (BHI) agar. After overnight growth we recovered the biofilm and enumerated colony forming units (CFU) of the different cell types (passage 0 in Fig. 1C and 1D). We observed a significant reduction in the number of erythromycin- and tetracycline-resistant transconjugants only when recipient cells received a pKH88 derivative encoding a cognate CRISPR guide (Fig. 1C and 1D). Non-cognate guides were ineffective at reducing antibiotic resistance. This loss of antibiotic resistance was maintained after growth of the community in fresh BHI broth overnight for the pKH88[sp-*ermB*] targeting group (passage 1 in Fig. 1C), and trended toward maintenance of antibiotic resistance loss for the pKH88[sp-*tetM*] targeting group (passage 1 in Fig. 1D).

### Conjugation of pKH88[sp-*ermB*] to MDR *E. faecalis* resolves erythromycin resistance

To determine whether the pKH88 plasmids would be effective against MDR *E. faecalis*, we used pKH88[sp-*ermB*] to target *ermB* in *E. faecalis* V583 (Fig. 2A), a vancomycin-resistant hospital-adapted isolate (44). V583 *ermB* is encoded on the plasmid pTEF1, and V583 does not encode a *tetM* gene (10, 45). Initially, we observed poor conjugation frequencies of pKH88 derivatives into V583 when using CK135 as the donor (Fig. S2A and S2B). *E. faecalis* V583 encodes a Type IV restriction endonuclease that restricts entry of 5-methylcytosine-modified DNA and has been characterized as a barrier to DNA uptake in enterococci (40, 46, 47). We reasoned that this barrier could account for the reduced dissemination of pKH88 to *E. faecalis* V583. Therefore, we utilized the *E. faecalis* OG1 variant strain OG1RF deleted for a 5-methylcytosine DNA methyltransferase (ΔEfaRFI) as the donor strain. This strain cannot methylate plasmid DNA and the plasmid will not be cleaved by the recipient cell following conjugation (47). We observed that delivery of pKH88[sp-*ermB*] from *E. faecalis* OG1RF(ΔEfaRFI) to *E. faecalis* V583 decreased total erythromycin resistance in the population (Fig. 2B). In addition, *E. faecalis* V583 did not tolerate CRISPR cleavage of *ermB* on pTEF1, indicated by a reduction in viable *E. faecalis* V583 cells (Fig. 2B) and which is consistent with our previous findings (26). This was specific to *ermB* cleavage, as transconjugants that received the nonspecific targeting plasmid pKH88[sp-*tetM*] retained cell viability and erythromycin resistance (Fig. 2C).

**Figure 2.**
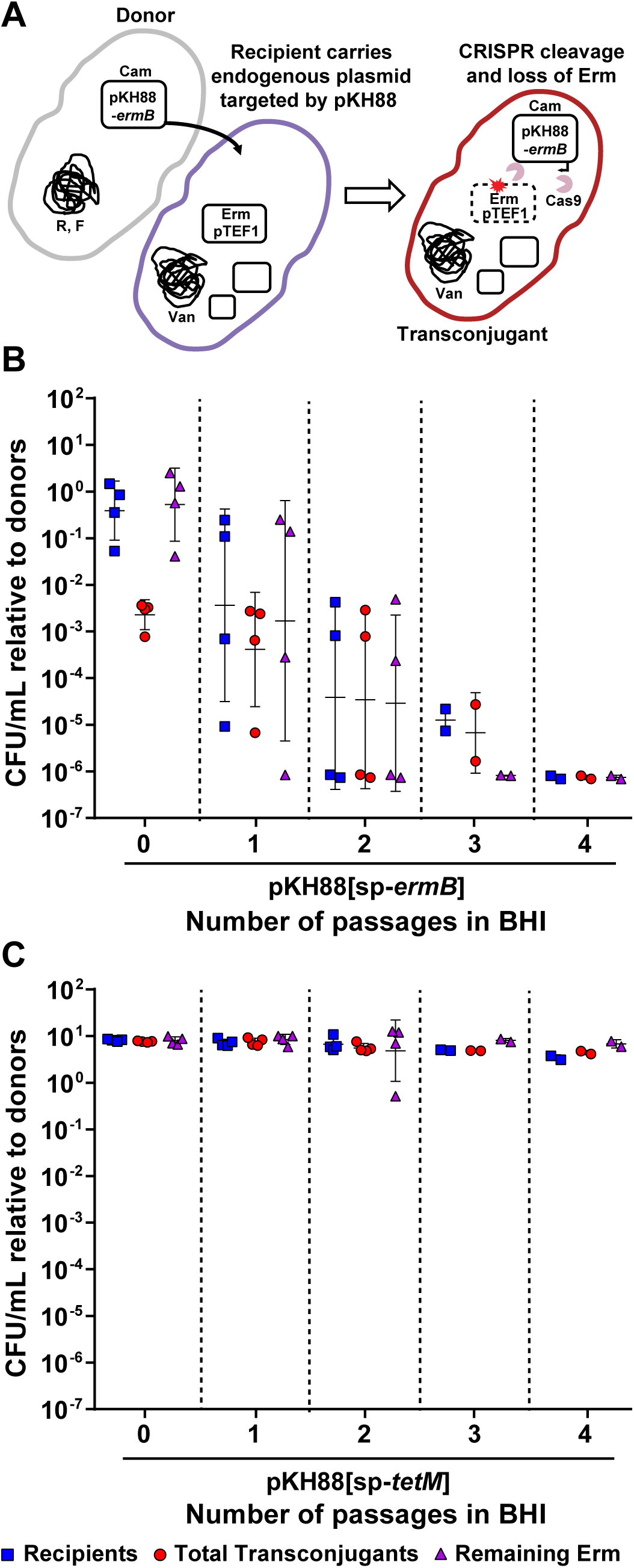
Engineered CRISPR-Cas targeting plasmid specifically reduces drug resistance in MDR *E. faecalis*. **(A)** Cartoon depicting the transfer, targeting and removal of erythromycin resistance from MDR *E. faecalis* V583. Conjugation of *E. faecalis* V583 with pKH88[sp-*ermB*] **(B)** but not pKH88[sp-*tetM*] **(C)** reduces erythromycin resistant *E. faecalis* V583 over the course of four serial passages in BHI broth. Markers; Erm – erythromycin, Van – vancomycin, R – rifampicin, F – fusidic acid, S – streptomycin, Sp – spectinomycin, Cam – chloramphenicol.

### CRISPR-Cas9 selectively depletes erythromycin resistance within enterococcal populations in the dysbiotic murine intestine

We next evaluated the efficacy of pKH88[sp-*ermB*] for the removal of erythromycin resistance using a mouse model of *E. faecalis* intestinal colonization. Mice were administered an antibiotic cocktail in their drinking water for seven days. Antibiotic water was removed for 24 hours and the mice were gavaged with *E. faecalis* OG1SSp(pAM771) recipients, modeling a bloom of antibiotic-resistant *E. faecalis* in the intestine. After an additional 24 hours, mice were gavaged with donor *E. faecalis* CK135 carrying either pKH88[sp-*ermB*] (targeting) or pKH88[sp-*tetM*] (non-targeting), and fecal samples were collected over the course of 23 days. A schematic of this experiment is depicted in Fig. S3. Whereas *E. faecalis* CK135 donors colonized the intestine to a sustained high density, *E. faecalis* OG1SSp(pAM771) recipient cells gradually declined in abundance over time (Fig. S4A and S4B). We attribute this gradual reduction in the recipient population to competitive killing by bacteriocin-producing donors as previously described for pPD1 carrying *E. faecalis* (29), and/or competition from the reemerging native microbiota that was initially displaced by the antibiotic treatment. Robust numbers of pKH88-containing transconjugants (i.e. chloramphenicol resistant OG1SSp) appeared two days post-colonization of the donor strains for both the pKH88[sp-*ermB*] and pKH88[sp-*tetM*] groups (Fig. S4C). However, the initial frequency of transconjugants per recipient reached only ∼10^-5^, indicating that overall conjugation frequency was relatively low (Fig. 3A). This was followed by a gradual increase in the transconjugant to recipient ratio, approaching 1:1 (Fig. 3A), as the total recipient population decreased over time (Fig. S4A). This suggests that the initial transconjugant population remains fixed due to a pKH88 bacteriocin-mediated fitness advantage upon restoration of the indigenous microbiota.

**Figure 3.**
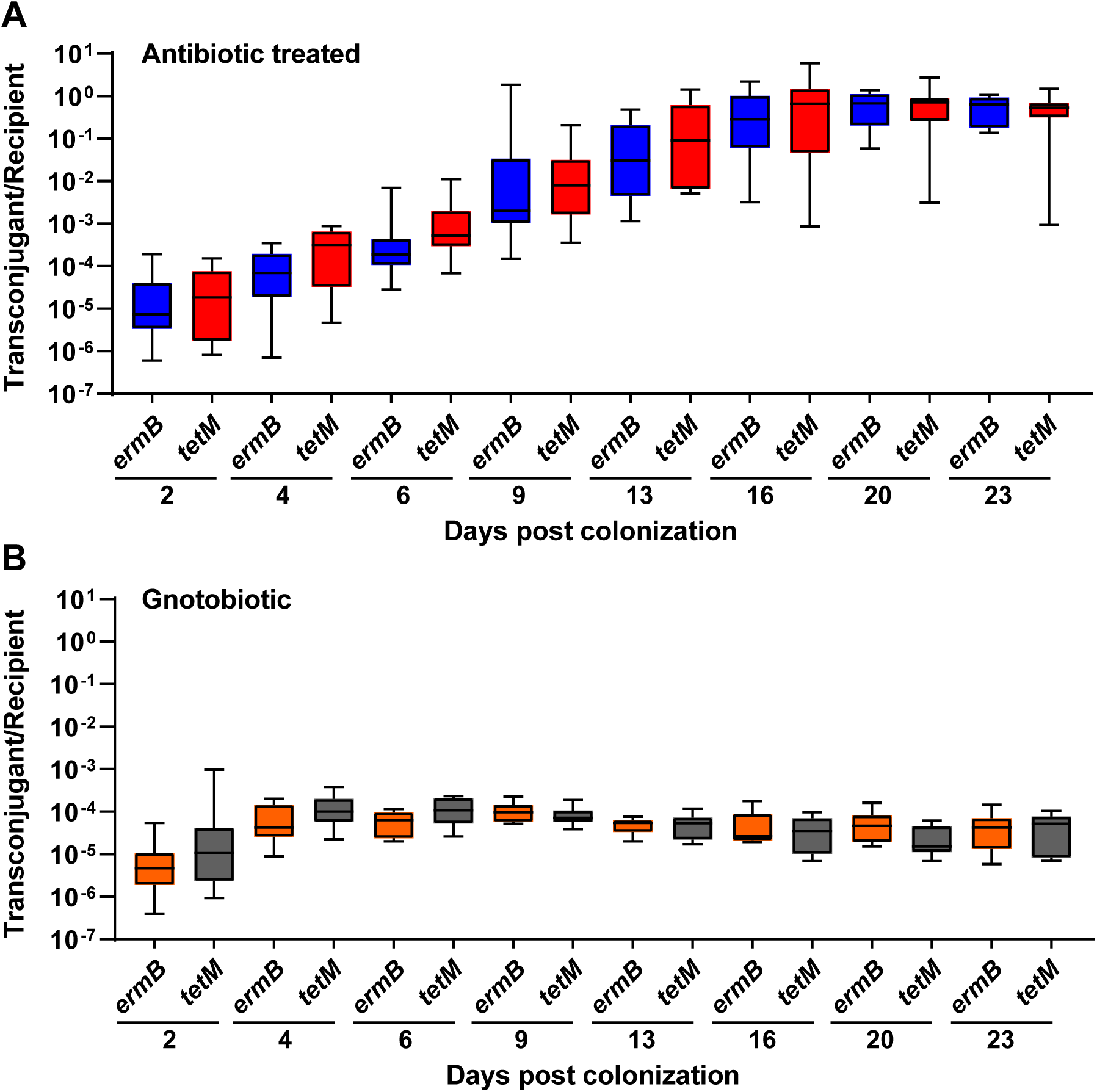
Saturation of the *E. faecalis* OG1SSp(pAM771) recipient population by pKH88 is time-dependent. The ratio of transconjugants to recipients was calculated for both the pKH88[sp-*ermB*] targeting group (*ermB*) and the pKH88[sp-*tetM*] non-targeting group (*tetM*) for 23 days of co-colonization in antibiotic-treated **(A)** and gnotobiotic **(B)** mice.

Although there appeared to be a trend of reduced erythromycin resistance within the recipient population for the pKH88[sp-*ermB*] group, no statistically significant reduction in the number of erythromycin-resistant recipients compared to the total recipient population was observed in the pKH88[sp-*ermB*] targeting group (Fig. 4A). Similarly no significant difference in the erythromycin resistant recipients compared to total recipients was observed for the pKH88[sp-*tetM*] non-targeting group (Fig. 4B). However, to our surprise, the difference in erythromycin-resistant recipients between the pKH88[sp-*ermB*] and pKH88[sp-*tetM*] groups was considerably less than the difference between the total erythromycin resistant intestinal *E. faecalis,* which includes both donors and recipients. The magnitude of this difference was ∼3 log by the end of the experiment (Fig 5A). This indicated that there was likely a substantial amount of conjugation by pAM771 into the donor population. Indeed, in as little as two days, donors acquired erythromycin resistance, but only in the pKH88[sp-*tetM*] non-targeting group (Fig. 5B). These data indicate that *E. faecalis* CK135(pKH88[sp-*tetM*]) donors are susceptible to counter-conjugation of pAM771 from recipients but that CK135(pKH88[sp-*ermB*]) donors are immune, presumably due to CRISPR-Cas targeting of *ermB* on pAM771 upon counter-conjugation. We conclude that our CRISPR antimicrobial dramatically reduces the prevalence of a targeted conjugative plasmid within the input *E. faecalis* population, likely through a multifactorial mechanism involving a bacteriocin-dependent competitive advantage, cleavage of the targeted plasmid, and prevention of counter-plasmid acquisition. This in turn substantially impacts the distribution of drug resistance, as mice colonized with pKH88[sp-*ermB*] donors had 0.1% of the total levels of erythromycin-resistant *E. faecalis* possessed by mice colonized with pKH88[sp-*tetM*] donors.

**Figure 4.**
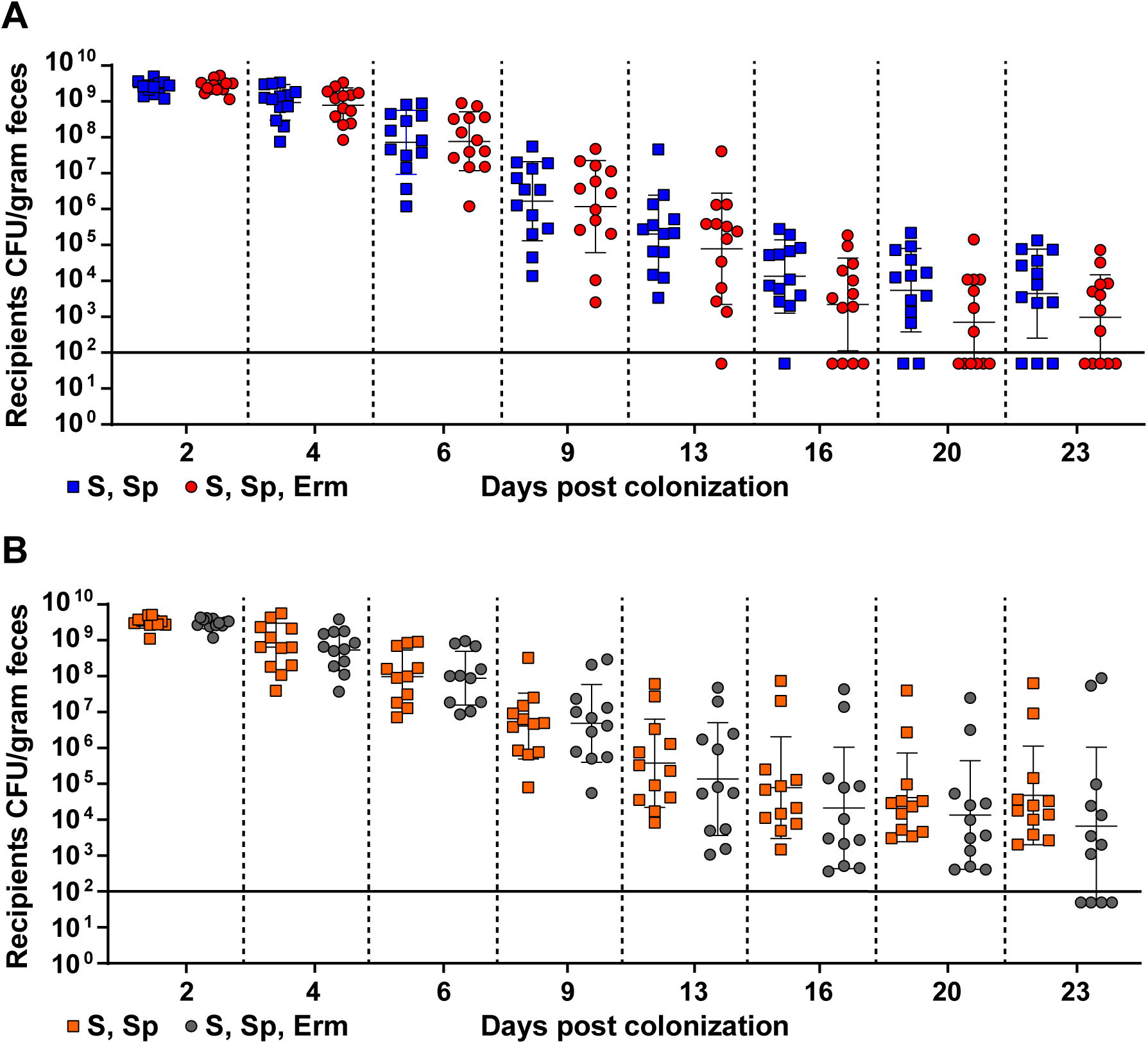
Assessment of erythromycin-resistant transconjugants from pKH88[sp-*ermB*] targeting and pKH88[sp-*tetM*] non-targeting populations in the murine intestine. **(A)** Antibiotic treated mice co-colonized with *E. faecalis* OG1SSp(pAM771) and CK135(pKH88[sp-*ermB]*) – targeting group. **(B)** Antibiotic treated mice co-colonized with *E. faecalis* OG1SSp(pAM771) and CK135(pKH88[sp-*tetM*]) – non-targeting group. The total OG1SSp(pAM771) recipient population and recipients that are erythromycin resistant were enumerated by selective plating of mouse feces over the course or 23 days of co-colonization. The solid horizontal line indicates the limit of detection. Markers; S – streptomycin, Sp – spectinomycin, Erm – erythromycin resistance.

**Figure 5.**
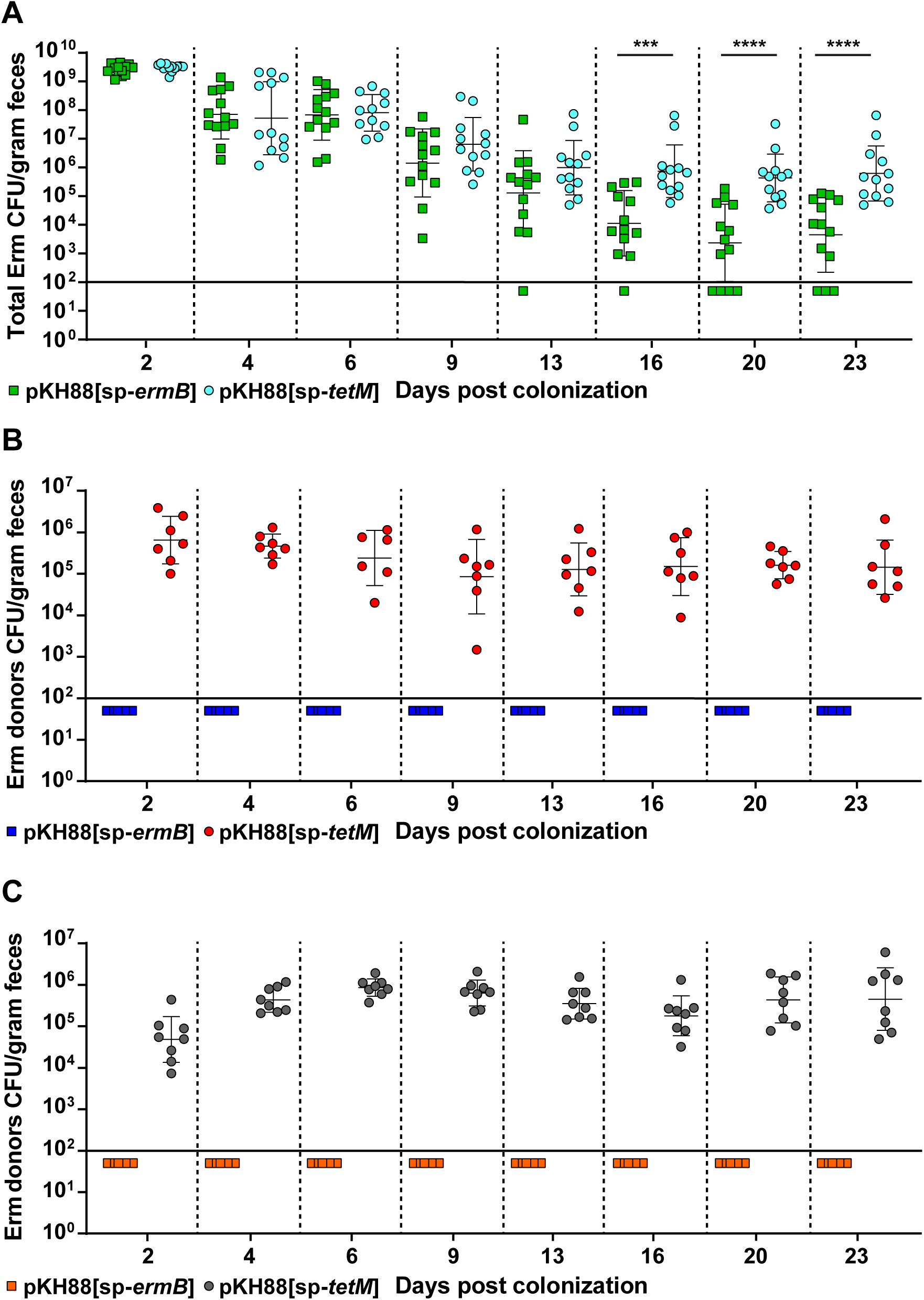
*E. faecalis* CK135 pKH88[sp-*ermB*] donors are immune to acquisition of erythromycin resistance from recipients in the murine intestine. **(A)** Total intestinal load of erythromycin resistance among input *E. faecalis* donors and recipients from antibiotic-treated mice. Erythromycin resistance is acquired by donors only in the pKH88[sp-*tetM*] non-targeting group in both antibiotic-treated **(B)** and gnotobiotic **(C)** mice. The solid horizontal line indicates the limit of detection. Markers; Erm – erythromycin. ***p=0.0003, ****p=<0.0001.

We next assessed whether the microbial complexity of the intestinal environment plays a role in the efficacy of the CRISPR-Cas antimicrobial, since curtailing antibiotic therapy spontaneously restores the microbiota in the antibiotic dysbiosis model (48). We first colonized germ-free mice with *E. faecalis* OG1SSp(pAM771) recipient cells and 24 hours later introduced donor *E. faecalis* CK135 carrying either pKH88[sp-*ermB*] or pKH88[sp-*tetM*]. Gnotobiotic mice maintained a high intestinal burden of both donors and recipients for the entire 23 days, in contrast to the antibiotic dysbiosis model where the recipient population is reduced over time (Fig. S5A, S5B and S4A). Despite sustained high density colonization of both donors and recipients, the total number of transconjugants was similar to those observed in the antibiotic dysbiosis model (Fig. S5C and S4C) but the transconjugant per recipient ratio remained stably low at 10^-5^ (Fig. 3B). Sustained high recipient density indicated that the apparent competitive advantage possessed by pKH88-containing donors via bac-21 does not manifest in gnotobiotic mice. These data suggest that even with prolonged high densities of both donors and recipients in the absence of microbial complexity, there may be biogeographical barriers that restrict the dissemination of pKH88 derivatives. This also implies that the competitive advantage possessed by the donor is most prevalent in the presence of the microbiota, and therefore some degree of microbial complexity is likely necessary for the functionality of our CRISPR-Cas antimicrobial. Interestingly, similar to observations in the antibiotic dysbiosis model, pKH88 donors acquired erythromycin resistance, but only in the pKH88[sp-*tetM*] non-targeting group (Fig. 5C).

These results demonstrate that our engineered CRISPR-Cas antimicrobial is a robust barrier to *in vivo* plasmid acquisition and reduces the *in vivo* prevalence of a specifically targeted drug resistance trait. From a technological viewpoint this is the ideal situation of a prospective “probiotic” donor; a strain that can effectively disseminate a CRISPR targeting plasmid while being protected from acquisition of undesirable traits.

### CRISPR-Cas antimicrobial prevents antibiotic-mediated intestinal expansion of transconjugants

A hallmark of enterococcal intestinal colonization is its ability to expand its population upon antibiotic perturbation of the indigenous microbiota (4, 25). This antibiotic-mediated expansion is attributed to both intrinsic and acquired antibiotic resistance. To determine the ability of our CRISPR-Cas antimicrobial to minimize the outgrowth of erythromycin resistant *E. faecalis* in the intestine following antibiotic therapy, we utilized both our antibiotic dysbiosis and gnotobiotic mouse models. We treated the mice 27 days post co-colonization with a single 40 µg dose of oral erythromycin and determined the abundance of erythromycin resistant transconjugants and total erythromycin resistant *E. faecalis*. Prior to antibiotic treatment there were no erythromycin resistant transconjugants recovered from mice receiving donors carrying the targeting plasmid pKH88[sp-*ermB*], whereas two of eight mice receiving donors carrying the non-targeting plasmid pKH88[sp-*tetM*] had recoverable erythromycin resistant transconjugants (Fig. 6). Upon oral erythromycin treatment, there was a ∼4 log expansion of erythromycin resistant transconjugants in the pKH88[sp-*tetM*] non-targeting group, and no erythromycin resistant transconjugants emerged in the mice colonized with *E. faecalis* CK135 pKH88[sp-*ermB*] donors (Fig. 6). This suggests that the perceived absence of erythromycin resistant transconjugants in six of the eight mice in the pKH88[sp-*tetM*] non-targeting group was due to their abundance being below the limit of detection prior to antibiotic treatment. Additionally, the total number of erythromycin-resistant *E. faecalis* in the pKH88[sp-*tetM*] non-targeting group was significantly higher compared to the pKH88[sp-*ermB*] targeting group (Fig. 6). Together these data indicate that transconjugants receiving pKH88[sp-*ermB*] have been depleted of erythromycin resistance and that pKH88[sp-*ermB*] can significantly reduce the overall total erythromycin resistance within intestinal *E. faecalis*, pre- and post-antibiotic therapy. Interestingly, any remaining viable transconjugants in the pKH88[sp-*ermB*] targeting group were impaired in their ability to increase in cell density compared to mice receiving pKH88[sp-*tetM*] non-targeting donors (Fig. 6), further indicating that targeted *E. faecalis* cells fail to expand their population due to loss of pAM771 encoded erythromycin resistance.

**Figure 6.**
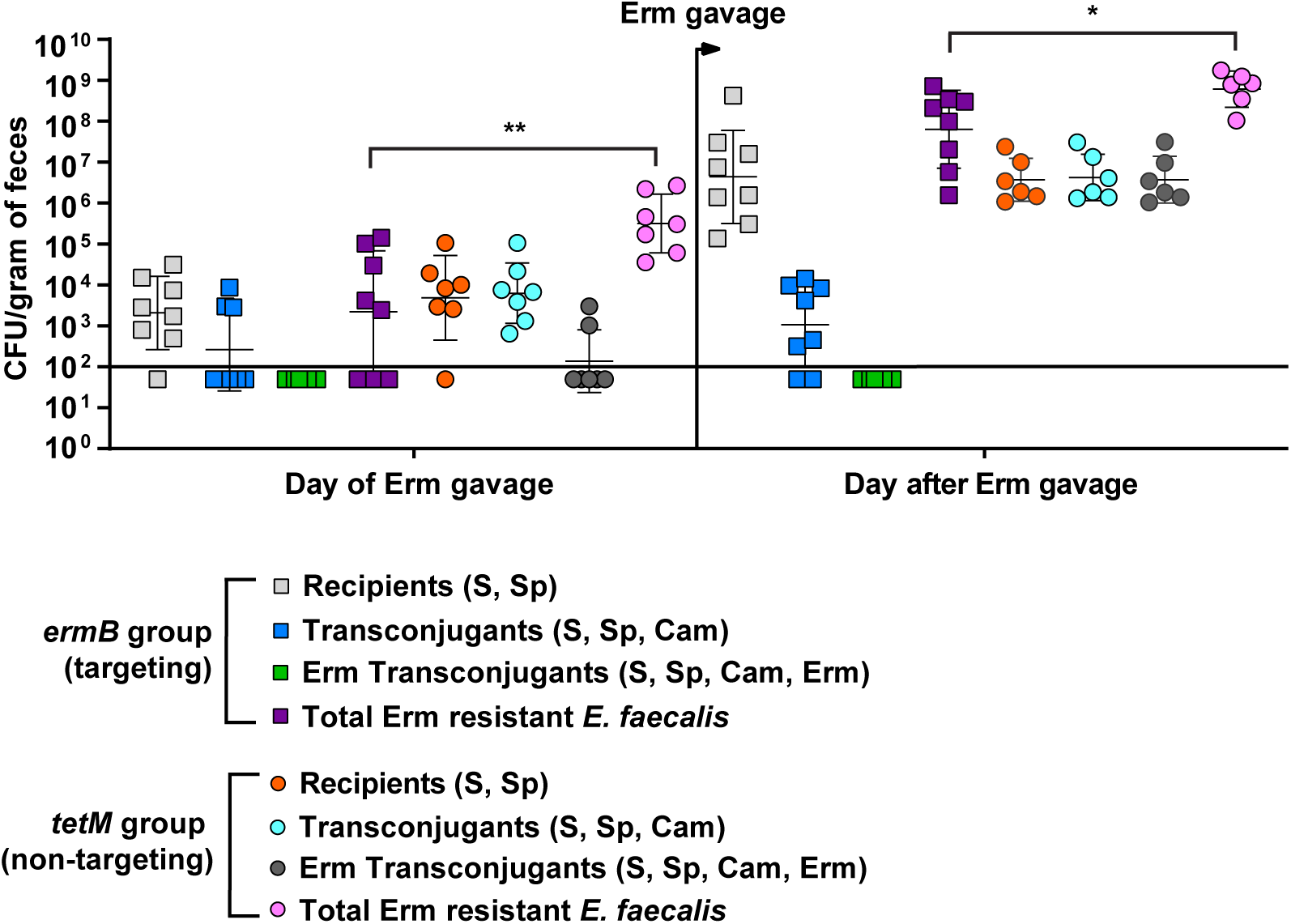
Transconjugants that receive pKH88[sp-*ermB*] do not bloom in the intestines of dysbiotic mice following oral erythromycin treatment. 27 days post co-colonization with *E. faecalis* OG1SSp(pAM771) and CK135(pKH88[sp-*ermB*]) (targeting group) or with *E. faecalis* OG1SSp(pAM771) and CK135(pKH88[sp-*tetM*]) (non-targeting group), mice received a single 40 µg dose of oral erythromycin. The number of recipients, transconjugants and erythromycin resistant transconjugants were enumerated from fecal pellets before and after oral erythromycin treatment. The solid horizontal line indicates the limit of detection. Markers; Erm – erythromycin, S – streptomycin, Sp – spectinomycin, Cam – chloramphenicol. *p=0.01, **p=0.004.

In gnotobiotic mice, following oral erythromycin treatment, again we did not recover any erythromycin resistant transconjugants in the pKH88[sp-*ermB*] targeting group (Fig. S6). Interestingly, in gnotobiotic mice, we did not observe outgrowth of erythromycin resistant transconjugants in the pKH88[sp-*tetM*] non-targeting group, even though we did not recover any erythromycin resistant transconjugants in the pKH88[sp-*ermB*] targeting group, similar to the antibiotic dysbiosis model (Fig. S6). We attribute this to the high level of both donors and recipients in the gnotobiotic intestine that were unaffected by oral erythromycin treatment (Fig. S5A and S5B). Intestinal colonization saturation by donors and recipients may be spatially restricting, thus promoting colonization resistance against transconjugant expansion upon erythromycin treatment in the pKH88[sp-*tetM*] non-targeting group.

## Discussion

This study presents a strategy for the delivery of a CRISPR-based antimicrobial that targets antibiotic-resistant *E. faecalis*. We show that this conjugative antimicrobial significantly reduces antibiotic resistance from *in vitro* and *in vivo* enterococcal communities in the absence of externally applied selection for donor strains. Our approach for the delivery of a CRISPR-Cas antimicrobial differs from studies which employed either transformable plasmids or phages for the recognition and delivery to target cells (18–21). Although these studies provided the foundation for the use of CRISPR antimicrobial activity against bacteria, limitations were noted. In the case of phage-mediated CRISPR antimicrobial dissemination, this technique suffered from incomplete phage dissemination. This scenario is expected due to selective pressures imposed on bacteria by phages during infection, including receptor mutations (22–25). Such issues have recently been addressed using multi-phage delivery systems or by co-opting phage particles for the delivery of CRISPR-Cas targeting modules embedded within mobilizable genome islands (49, 50). Although the use of a phage-based delivery system for *E. faecalis* should be possible, enterococcal phage biology is understudied and limited information exists concerning phage host ranges and genetics. Therefore, we chose to engineer a PRP for the dissemination of an *E. faecalis*-specific CRISPR-Cas antimicrobial for several reasons. First, the conditions governing PRP regulation have been extensively studied. Second, PRPs are specific for *E. faecalis*. Third, PRPs achieve high conjugation rates and can be genetically modified to diversify host tropism by altering pheromone responsiveness. Fourth, PRPs are stably maintained and disseminate in the absence of selection (51, 52).

The intestine is a reservoir for MDR *E. faecalis*, and *E. faecalis* infections often follow exposure to broad spectrum antibiotics. We applied our CRISPR-Cas antimicrobial method for the removal of an antibiotic resistance trait from intestinal *E. faecalis*. The mouse models of intestinal colonization used were designed to mimic enterococcal overgrowth in the human intestine following antibiotic therapy (3, 4). Antibiotics disrupt the intestinal microbiota, allowing antibiotic-resistant enterococci to become the dominant colonizer. In our study, we used *E. faecalis* donor strains for the conjugative delivery of CRISPR-Cas to remove erythromycin resistance from an abundant intestinal *E. faecalis* target population. The conjugative transfer of an *ermB*-specific CRISPR-Cas targeting system into recipient *E. faecalis* cells showed little to no reduction in erythromycin resistance from the total intestinal *E. faecalis* recipient population. However, within the recipient population that did receive the CRISPR targeting plasmid carrying a cognate spacer, all transconjugants were fully depleted of erythromycin resistance. This indicates that, although the frequency of conjugation was low compared to the total available recipient population and not as robust as *in vitro* conjugation frequencies, the specificity of our CRIPSR-Cas targeting system in the intestine was absolute. If successful conjugation occurs, there is complete loss of erythromycin resistance. In addition, we observed that donors carrying a cognate CRISPR antimicrobial plasmid were protected from acquisition of erythromycin resistance from recipients. This is an important feature of our deliverable system since the transmission of undesirable target traits to donor cells would impede the usefulness of such a delivery strategy within complex microbial environments. The discovery that *E. faecalis* donor cells carrying the CRISPR-Cas antimicrobial are immunized against the acquisition of antibiotic resistance suggests that donors could be used as a probiotic. These probiotic *E. faecalis* cells would be protected from the acquisition of antibiotic resistance traits and could pre-emptively occupy the intestine to outcompete or inhibit invading multidrug resistant strains.

There are several reasons why intestinal conjugation frequency might be suboptimal in our study. First, our analysis measured transconjugants from fecal samples, not the intestinal mucosa. The spatial distribution of donor and recipient *E. faecalis* cells colonizing the intestinal mucosal layer is unknown and evidence suggests that conjugation frequency could be impacted by the regional location of *E. faecalis* intestinal colonization (28). Second, pKH88 derivatives require recipient-derived cPD1 pheromone to induce conjugation (53). We know little about the kinetics of pheromone signaling in the intestine. Low rates of conjugation could reflect low local concentrations of cPD1 in the vicinity of donors resulting in few donors that effectively conjugate, thereby minimizing conjugation frequencies. Alternatively, excess donors can inhibit conjugation to recipient cells (54). This is due to the production of inhibitory self-signaling peptides produced by the donors that compete with recipient pheromones and reduce conjugation frequencies. Exploring the efficiency of conjugation upon altering donor to recipient ratios could lead to more permissive conjugation in the intestine. Another possibility is that conjugation in the intestine may be more efficient at a precise density of donors, recipients, or both, which may require future optimization for effective plasmid delivery.

Moving forward, improvements to our current system are justified if an engineered conjugative CRISPR antimicrobial is to be clinically effective for precision removal of antibiotic resistance genes in *E. faecalis* or other MDR bacteria. Modifications include multiplexing CRISPR guide RNAs to target different resistance genes and/or the same genes at multiple sites, as well as increasing the overall conjugation frequency by altering PRP regulation, dosing of donor strains, or the plasmid replicon itself. Theoretically, resistance to bac-21 could compromise the efficacy of our system, and future iterations should utilize alternative bacteriocins as well as bacteriocin-free constructs. In this study we focused on targeting antibiotic resistance that resides on plasmids. However, many *E. faecalis* antibiotic resistance genes are chromosomally encoded. Future iterations of this system should also be engineered for the specific targeting of chromosomal antibiotic resistance genes. Recently we showed that CRISPR-Cas targeting of the *vanB* locus on the *E. faecalis* V583 chromosome is lethal (40). Targeting chromosomal antibiotic resistance genes could have the added feature of bacterial lethality. Such a strategy may be preferable depending on whether the intended use of the CRISPR-Cas antimicrobial is for reshaping the microbiota or for targeting drug resistant strains during infection.

Our model system has relied on laboratory-domesticated strains of *E. faecalis* OG1 (43), it will be imperative to determine the efficacy of this system against more “wild” clinical isolates which are the intended targets of such a therapy. For safe and effective delivery of the CRISPR antimicrobial in the clinic, the use of an attenuated *E. faecalis* strain or a related member of the Lactobacillales approved for probiotic use would be essential. Finally, this system could be adapted for the targeted removal of antibiotic resistance from bacteria other than *E. faecalis.* Gram-positive conjugative plasmids like pIP501, which belongs to the *rep*_1_ family, are broadly disseminated conjugative plasmids (51, 52, 55). pIP501 is transferrable among enterococci, other lactic acid bacteria, staphylococci, and clostridia (55). The modification of a promiscuous mobilizable plasmid such as pIP501 with CRISPR-Cas antimicrobial function could expand the capabilities of our current system by facilitating broad targeting of multiple Gram-positive pathogens.

## Materials and Methods

### Bacterial culture methods and molecular biology techniques

*Escherichia coli* cultures were incubated with shaking at 220 RPM at 37°C in Lysogeny Broth (LB). *E. faecalis* cultures were grown in Brain Heart Infusion (BHI) broth at 37°C without shaking. PCR was performed using Taq Polymerase (New England Biolabs), and Q5 DNA Polymerase (New England Biolabs) for molecular cloning. The PureLink PCR Purification kit (Invitrogen) was used to purify DNA fragments and the GeneJet Plasmid Purification kit (Fisher) was used for plasmid purification. Primers were obtained from Sigma Aldrich. Sanger sequencing was performed at the Massachusetts General Hospital DNA Core Facility. *E. coli* EC1000 was used for routine plasmid propagation (56). *E. faecalis* and *E. coli* competent cells were prepared as described previously (26). Genomic DNA was extracted using the MO BIO Microbial DNA Isolation Kit (Qiagen). Antibiotics were used at the following concentrations: chloramphenicol, 15 μg/ml; streptomycin, 500 μg/ml; spectinomycin, 500 μg/ml; vancomycin, 10 μg/ml; erythromycin, 50 μg/ml; rifampicin, 50 μg/ml; fusidic acid, 25 μg/ml; tetracycline, 10 μg/ml.

### Strain and plasmid construction

All strains used in this study can be found in Table S1. The donor strain used for competition experiments was a rifampicin and fusidic acid-resistant derivative of CK135 (57). All plasmids used in this study can be found in Table S2. Fragments obtained from the plasmids pHA101 (58), pPD1 (24), P*_bacA_*-pG19, pKH12 (26) and pGR *tetM*/*ermB* (40) were amplified using Q5 DNA polymerase and assembled using NEB HiFi DNA Assembly Master Mix (New England Biolabs). For simplicity, assembly of the CRISPR-targeting construct (P*_bacA_-cas9*-*cat*-guide RNA) was divided into two steps. First, P*_bacA_cas9* was introduced into pPD1 by homologous recombination. Then the *cat* gene from pKH12 and the P*_bacA_*-driven CRISPR guide RNA was integrated downstream of P*_bacA_-cas9*. Following the generation of this construct, we discovered that the first integration site obstructed the pheromone response, therefore the entire engineered region (P*_bacA_-cas9*-*cat*-guide RNA) was amplified as one fragment and integrated into native pPD1 between orfs *ppd4* and *ppd5* using the integration plasmid pCOP88 (Fig. S1). The modified pPD1 derivatives are designated as pKH88[sp-*tetM*] and pKH88[sp-*ermB*], which target *tetM* or *ermB* respectively. Primer sequences are listed in Table S3.

### *In vitro* competition assays

Overnight cultures of donors and recipients were pelleted, resuspended in fresh BHI broth, and cultured for 1.5 hours. Donors and recipients were mixed in a volume ratio of 1:9, and the mixtures were spread on BHI agar. Following overnight incubation, lawns were scraped into sterile PBS and serial dilutions (10 µl each) were spotted on BHI plates supplemented with antibiotics to enumerate colony forming units (CFU) of donors, recipients, and transconjugants as well as total CFU of tetracycline- and erythromycin-resistant cells. For individual passages, the recovered conjugation mixture was diluted 1:1000 in fresh BHI broth without selection and incubated overnight, after which the constituents of the mixture were again enumerated.

### Mouse models of *E. faecalis* colonization

All animal protocols were approved by the Institutional Animal Care and Use Committee of the University of Colorado School of Medicine (protocol number 00253).

For the antibiotic dysbiosis model, 7 days prior to colonization, 6-8 week old C57BL6/J mice were gavaged with 150 μL of an antibiotic cocktail (streptomycin 1 mg/mL, gentamicin 1 mg/mL, erythromycin 200 μg/mL) and given a water bottle *ad libitum* with the same antibiotic cocktail for 6 days following the initial gavage. 24 hours prior to colonization with recipient *E. faecalis*, antibiotic water was removed and replaced with standard sterile reverse osmosis water. Donor and recipient strains were cultured overnight in BHI and mice were gavaged with ∼10^9^ CFU in 100 μL of sterile PBS. All mice were first gavaged with *E. faecalis* OG1SSp(pAM771). After 24 hours, mice (n=13, 7 female (F) and 6 male (M)) were gavaged with CK135(pKH88[sp-*ermB*]) while the second group (n=12, 5 F and 7 M) were gavaged with CK135(pKH88[sp-*tetM*]). Fecal samples were collected at the designated time points, resuspended in 1 mL of sterile PBS, and plated on BHI agar supplemented with rifampicin, fusidic acid, erythromycin, chloramphenicol, streptomycin and spectinomycin in combinations that would select for the desired populations and using the concentrations described above. After 27 days, 8 mice (4 F and 4 M) from the pKH88[sp-*ermB*] targeting group and 7 mice (3 F and 4 M) from the pKH88[sp-*tetM*] non-targeting group were gavaged with erythromycin (40 μg). Fecal samples were collected from all mice pre-erythromycin gavage. Post-erythromycin gavage, fecal samples were collected from all 8 mice from the pKH88[sp-*ermB*] targeting group and from 6 of 7 mice from the pKH88[sp-*tetM*] non-targeting group due to the inability of a single mouse from this group to defecate.

All germ-free mice were gavaged with *E. faecalis* OG1SSp(pAM771. 24 hours later these gnotobiotic mice were gavaged with *E. faecalis* CK135(pKH88[sp-*ermB*]) (n=8, 4 F and 4 M) or *E. faecalis* CK135(pKH88[sp-*tetM*]) (n=8, 4 F and 4 M). Fecal samples were collected at the designated time points and donors, recipients, and transconjugants were enumerated on selective media as described above. After 27 days, all 8 gnotobiotic mice from each group were gavaged with 40 μg of erythromycin.

### Statistical analysis

GraphPad-Prism version 8.2.0 was used to determine statistical significance and the following tests were employed based on the experimental set up. For *in vitro* competition assays a log normal distribution was assumed and a Student’s t-test was performed. For intestinal colonization studies, due to a higher degree of variation in the colonization kinetics between individual mice, a normal distribution was not assumed and thus a nonparametric Mann-Whitney test was performed. Values below the limit of detection for in vivo animal experiments were represented as constant (one half of the value of the limit of detection). The limit of detection for all animal experiments was 100 colony forming units per gram of feces. Error bars represent the geometric mean +/- the geometric standard deviation. p-values indicate the following: * 0.01 – 0.05, ** 0.001 - <0.01, *** 0.0001 - <0.001, **** <0.0001.

## Acknowledgements

This work was supported by R01AI116610 (KLP), R01AI141479 (BAD), and K01DK102436 (BAD). We thank members of the Palmer and Duerkop labs for critical feedback on the manuscript. We thank Christopher Kristich for providing *E. faecalis* CK135(pPD1).

**Figure S1.**
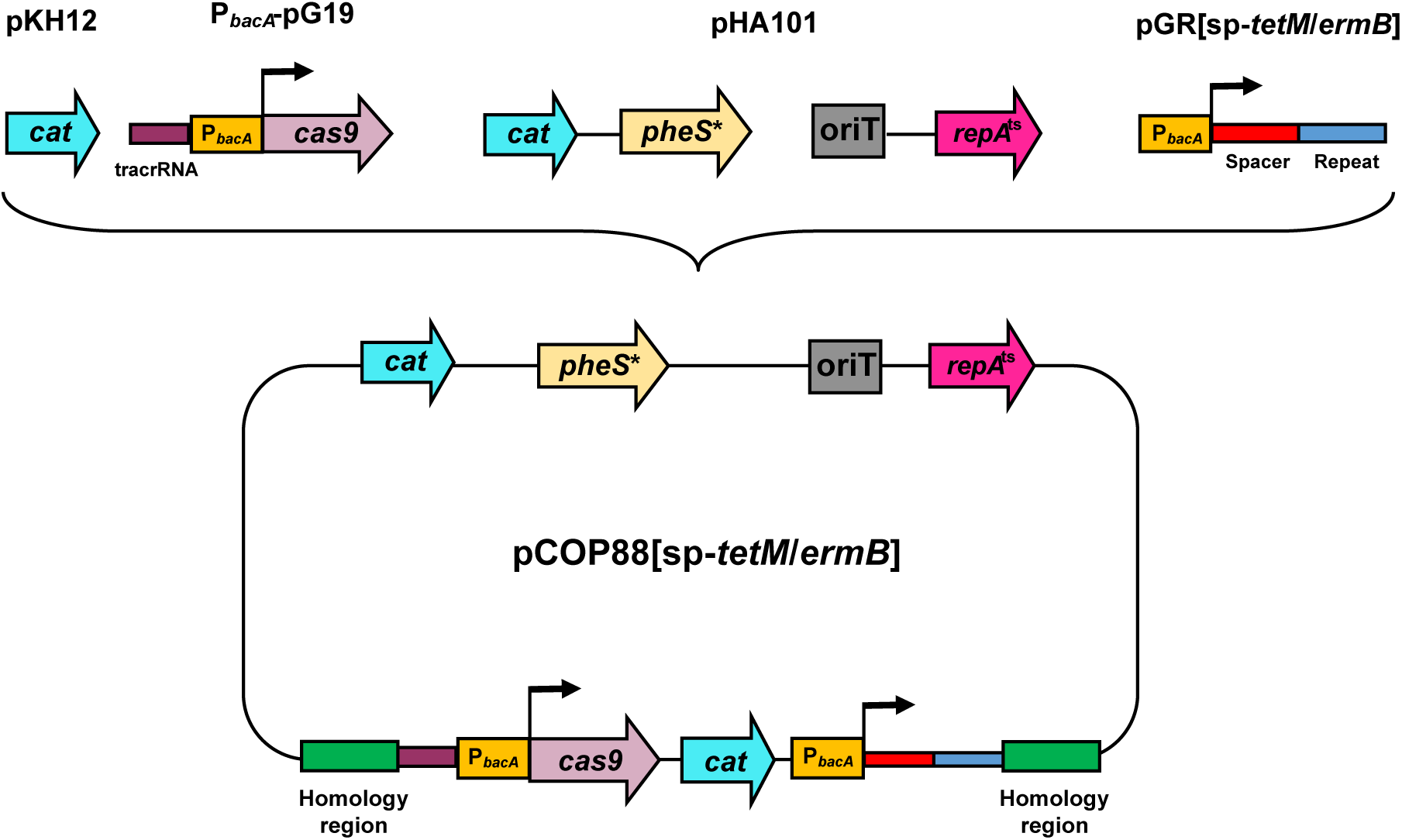
The allelic exchange vector pCOP88 integrates the P*_bacA_-cas9*-*cat*-guide RNA module into pPD1. pCOP88 derivatives contain elements from the allelic exchange vector pLT06 (59). These include a temperature sensitive *repA* (*repA*^ts^) allele that facilitates forced integration of pCOP88 derivatives into target DNA sequences under non-permissive temperature (42°C) and a P-*pheS* cassette that allows for the cellular utilization of *p*-chloro-phenylalanine as a counter-selectable marker. The P*bacA*-*cas9*-*cat*-guide RNA module is flanked by 1-kb pPD1 homology regions used for the homologous recombination of the module into native pPD1.

**Figure S2.**
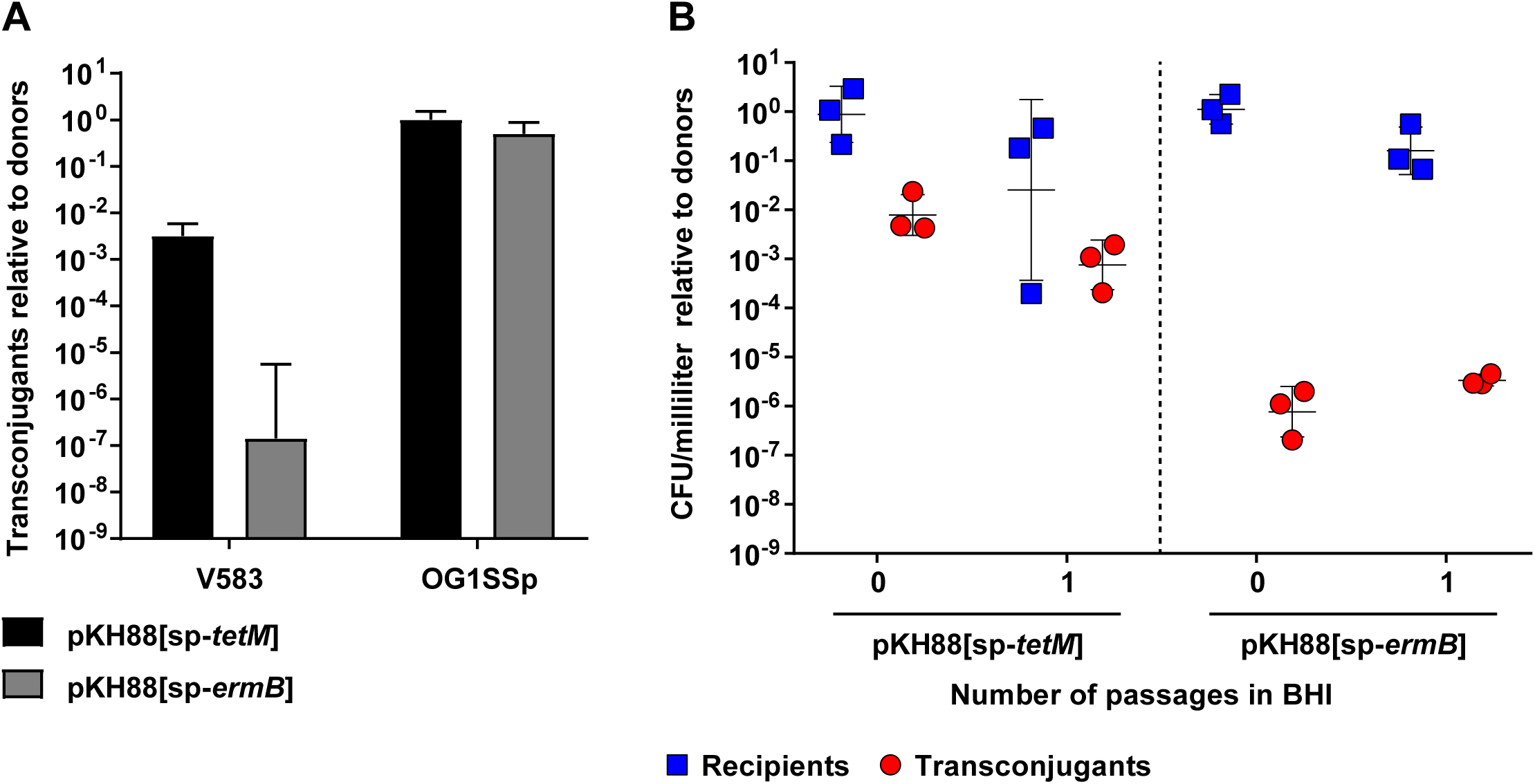
*E. faecalis* V583 restricts conjugation of pKH88 derivatives originating from *E. faecalis* CK135. **(A)** The ratio of *E. faecalis* V583 and OG1SSp transconjugants following *in vitro* co-culture with *E. faecalis* CK135(pKH88) derivative donors. **(B)** Comparison of normalized *E. faecalis* V583 transconjugant and *E. faecalis* V583 recipient numbers following *in vitro* co-culture with *E. faecalis* CK135(pKH88) derivative donors.

**Figure S3.**
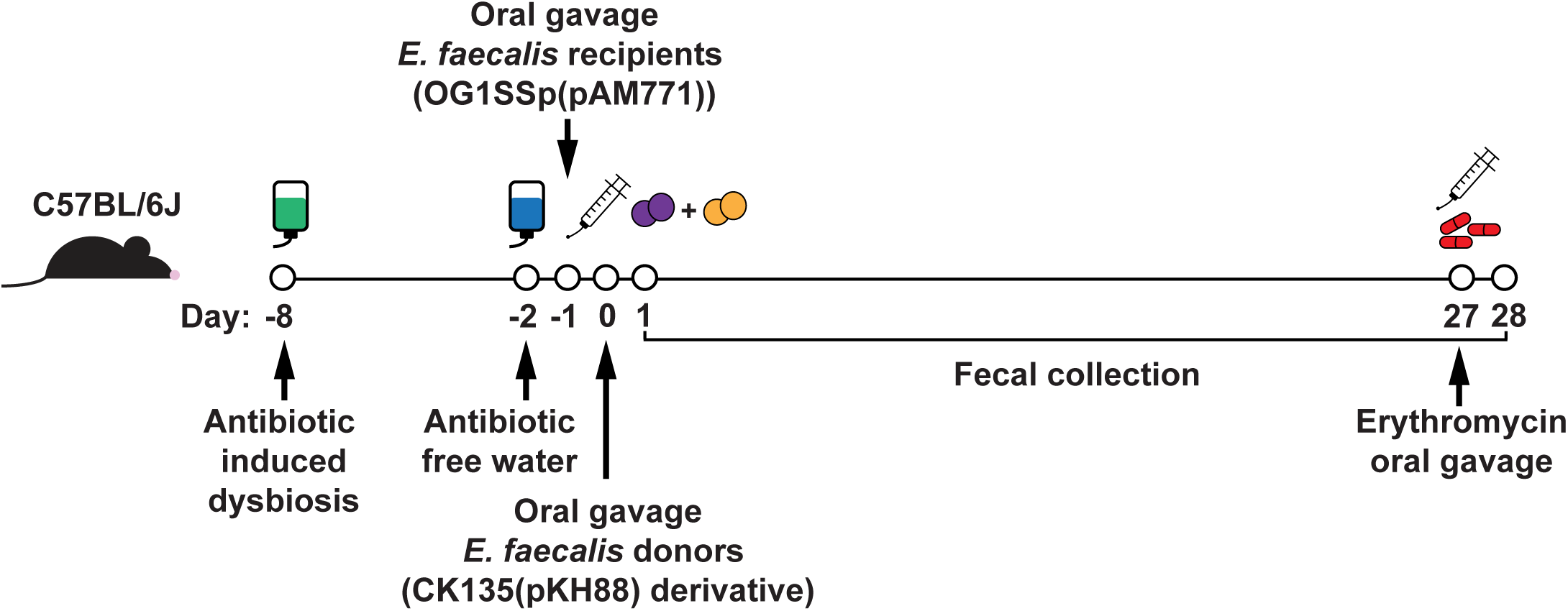
Schematic cartoon of the antibiotic dysbiosis mouse model.

**Figure S4.**
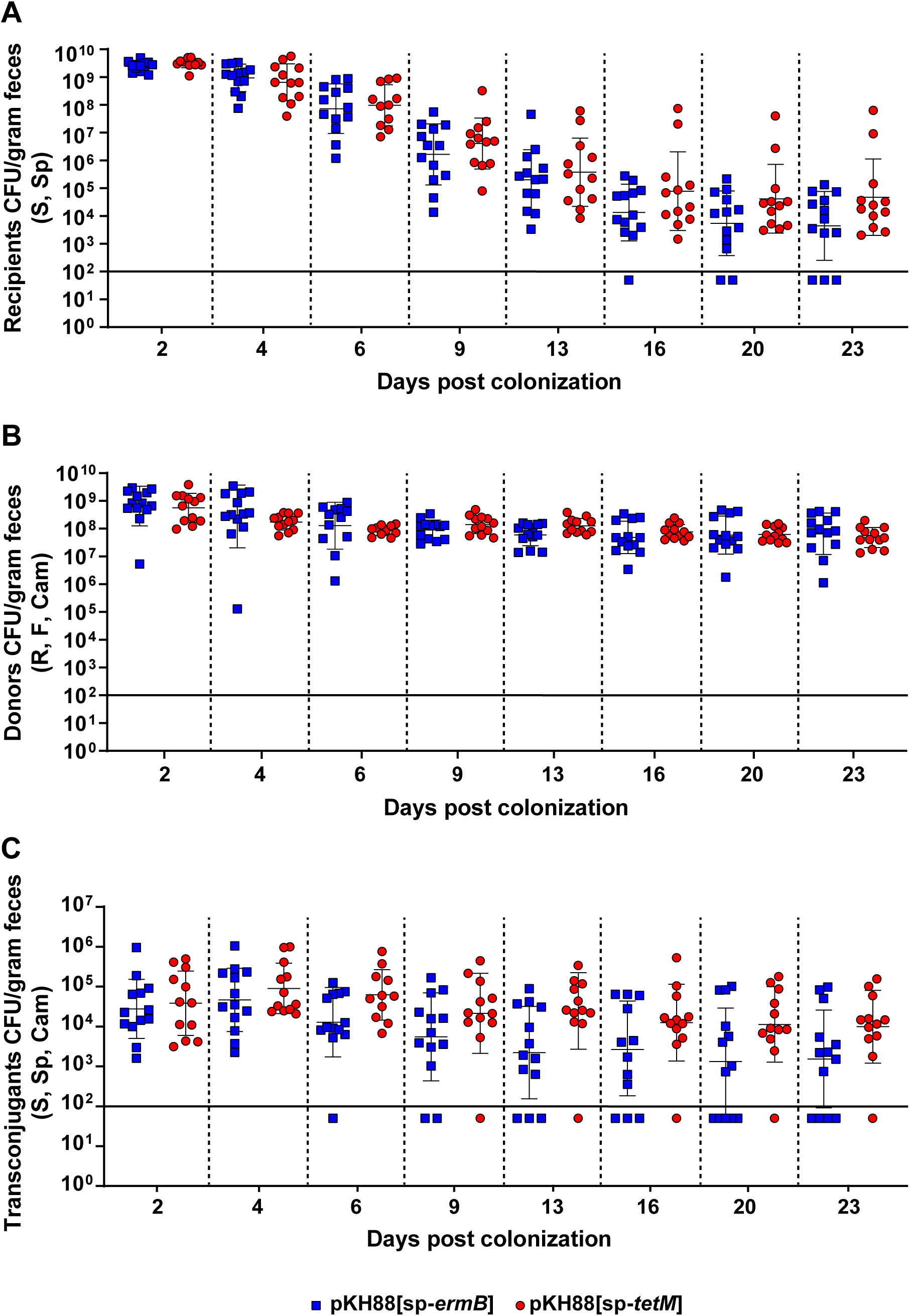
Intestinal colonization of *E. faecalis* populations following co-colonization with OG1SSp(pAM771) and CK135(pKH88) donors in antibiotic-treated mice. **(A)** Recipients. **(B)** Donors. **(C)** Transconjugants. Markers; S – streptomycin, Sp – spectinomycin, R – rifampicin, F – fusidic acid, Cam – chloramphenicol.

**Figure S5.**
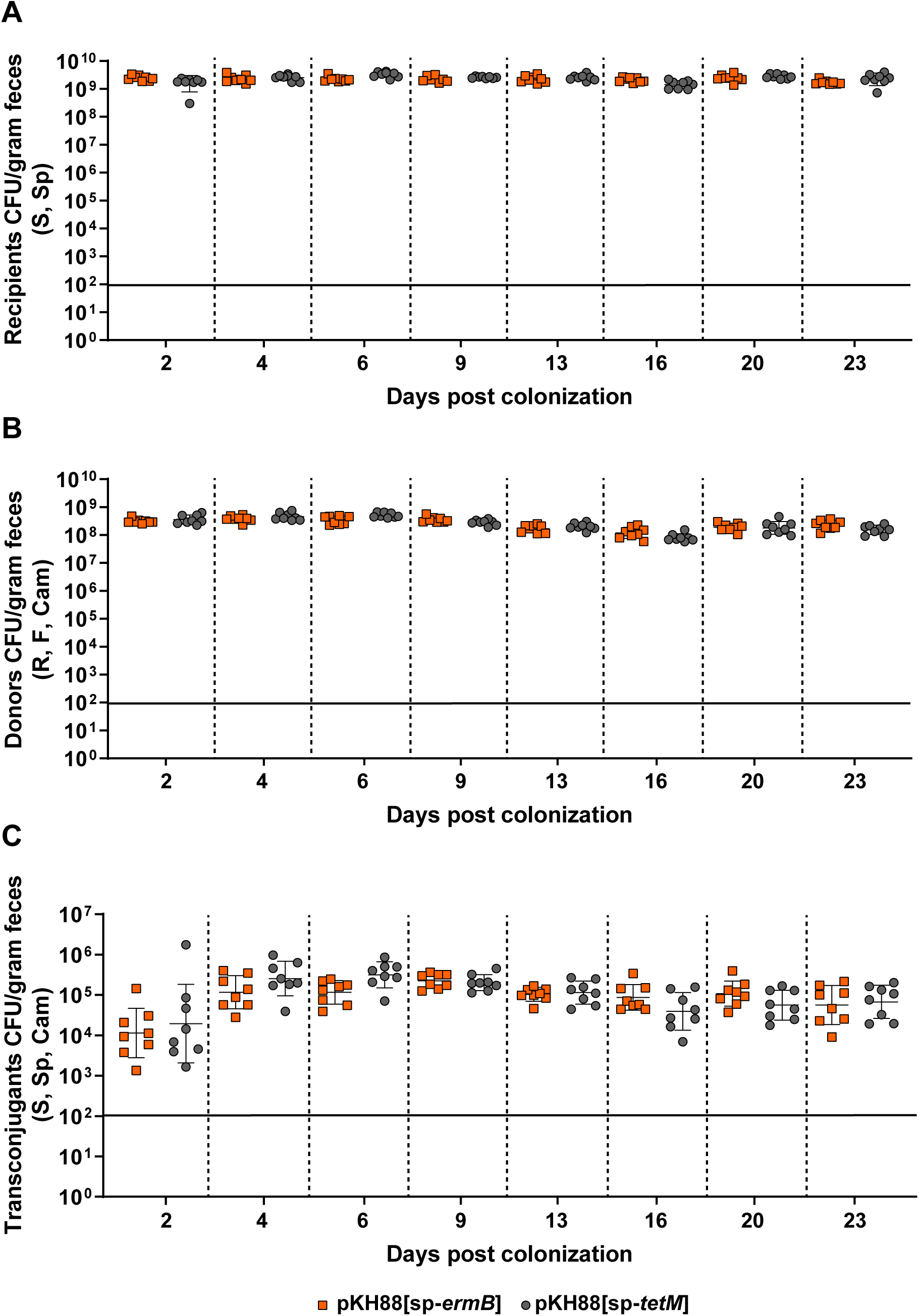
Intestinal colonization of *E. faecalis* populations following co-colonization with OG1SSp(pAM771) and CK135(pKH88) donors in gnotobiotic mice. **(A)** Recipients. **(B)** Donors. **(C)** Transconjugants. The solid horizontal line indicates the limit of detection. Markers; S – streptomycin, Sp – spectinomycin, R – rifampicin, F – fusidic acid, Cam – chloramphenicol.

**Figure S6.**
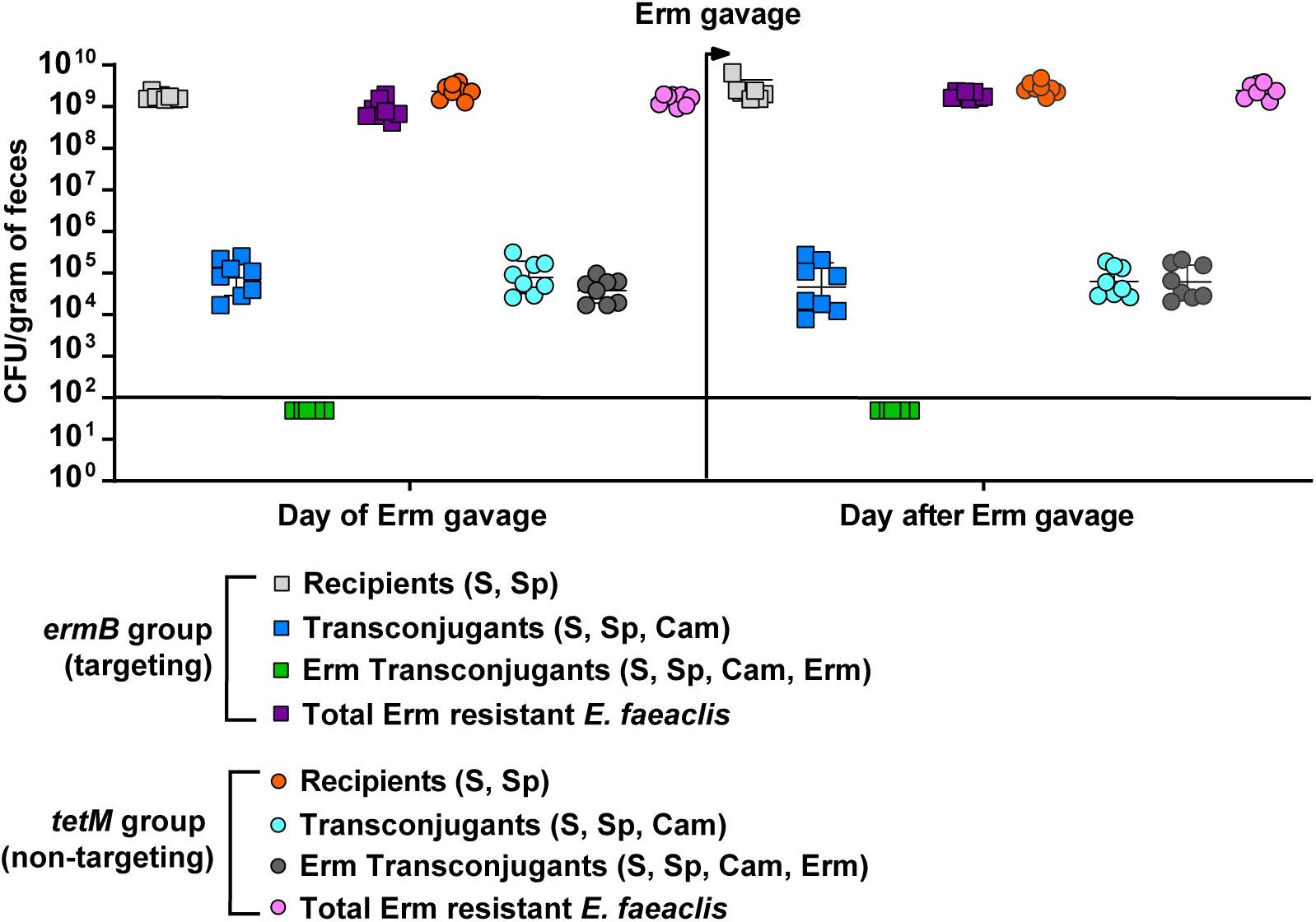
Non-targeted *E. faecalis* intestinal transconjugant populations do not bloom following oral erythromycin treatment of gnotobiotic mice. 27 days post co-colonization with *E. faecalis* OG1SSp(pAM771) and CK135(pKH88[sp-*ermB*]) (targeting group) or with *E. faecalis* OG1SSp(pAM771) and CK135(pKH88[sp-*tetM*]) (non-targeting group), mice received a single 40 µg dose of oral erythromycin. The number of recipients, transconjugants and erythromycin resistant transconjugants were enumerated from fecal pellets before and after oral erythromycin treatment. The solid horizontal line indicates the limit of detection. Markers; Erm – erythromycin, S – streptomycin, Sp – spectinomycin, Cam – chloramphenicol.

**Table S1.** Strains used in this study

| Organism | Strain Name | Description | Ref |
| --- | --- | --- | --- |
| *E. coli* | EC1000 | *E. coli* cloning host, providing *repA* in *trans*. F<sup>-</sup>, *araD139 (ara ABC-leu)*7679, *galU*, *galK*, *lacX74*, *rspL*, *thi*, *repA* of pWV01 in *glgB*, *km* | (1) |
| *E. faecalis* | V583 | MDR bloodstream isolate, resistant to vancomycin, gentamicin, and erythromycin | (2) |
|  | OG1SSp | Spectinomycin-streptomycin-resistant derivative of OG1 | (3) |
|  | CK135 | OG1 *rpoB*<sub>H486Y</sub> (spontaneous Rif<sup>r</sup> derivative) | (4) |
|  | CK135RF | Spontaneous fusidic acid-resistant derivative of CK135 | This study |
|  | OG1RF ΔEfaRFI | OG1RF EfaRFI (OG1RF_11622-11621) deletion mutant | (5) |

**Table S2.** Plasmids used in this study

| Plasmid | Description | Ref |
| --- | --- | --- |
| pLT06 | Encodes temperature-sensitive *repA* and *pheS* *<br>counter-selection | (6) |
| pHA101 | pLT06 + *oriT* | (7) |
| pCOP88[sp-*tetM*] | pLT06 derivative used to knock in CRISPR-targeting construct with a spacer targeting *tetM* | This study |
| pCOP88[sp-*ermB*] | pLT06 derivative used to knock in CRISPR-targeting construct with a spacer targeting *ermB* | This study |
| pKH88[sp-*ermB*] | pPD1 derivative with CRISPR-targeting cassette<br>for *ermB*; also encodes *cat* | This study |
| pKH88[sp-*tetM*] | pPD1 derivative with CRISPR-targeting cassette<br>for *tetM*; also encodes *cat* | This study |
| pAM771 | Non-cytolytic derivative of the PRP pAD1<br>mutagenized with Tn917, encodes erythromycin<br>resistance via *ermB* | (8) |
| pCF10 | PRP; encodes tetracycline resistance via *tetM* | (9) |
| pPD1 | PRP; encodes Bac-21 bacteriocin | (10) |

**Table S3.**
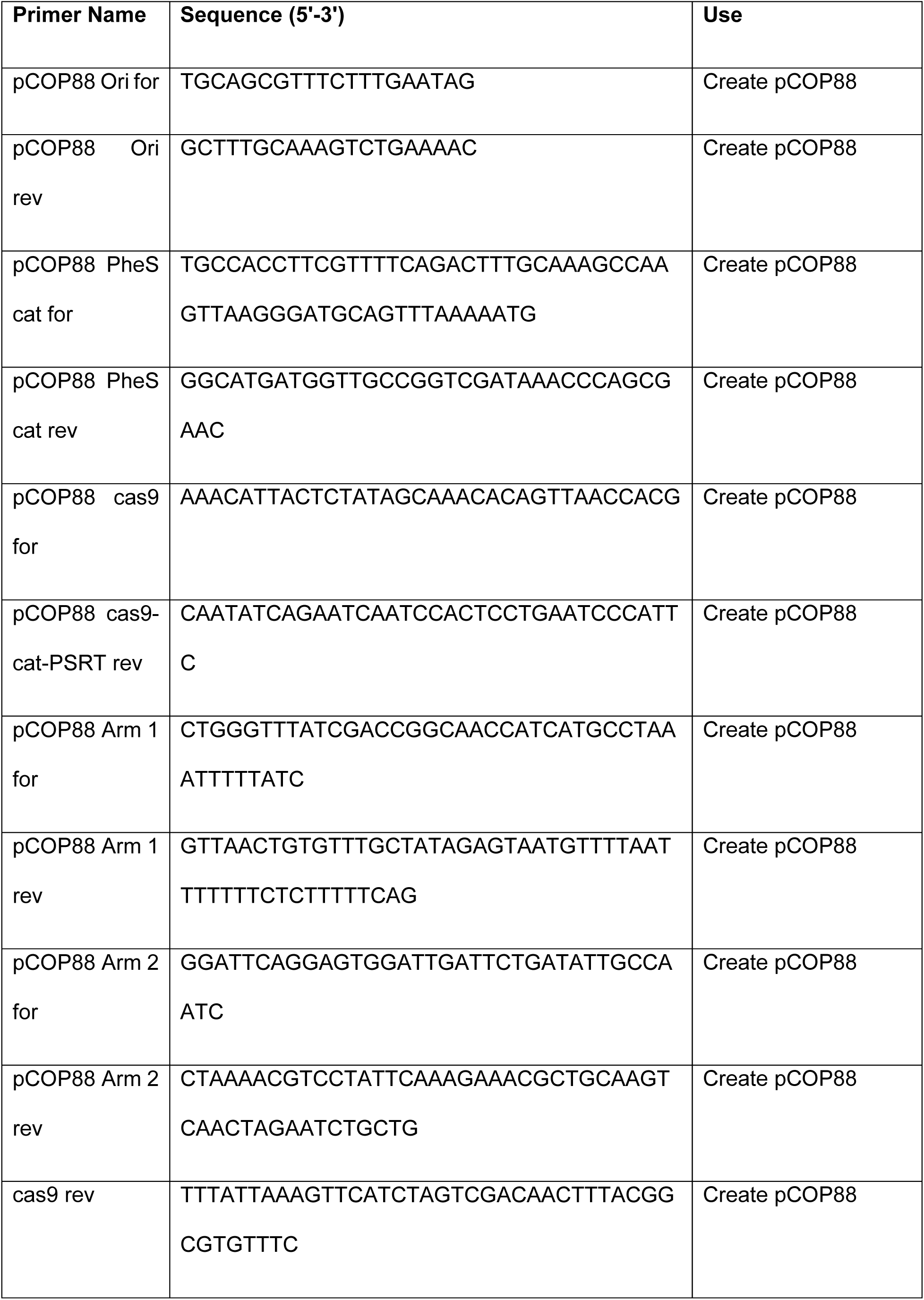
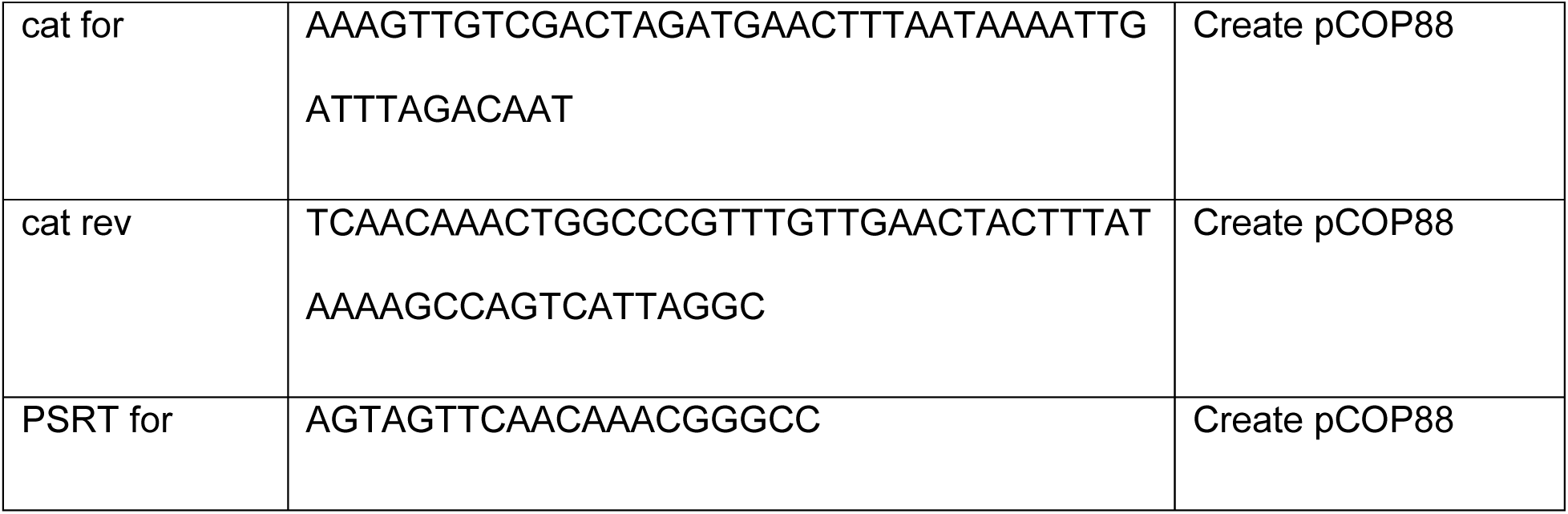
Primers used in this study

